# Distilling identifiable and interpretable dynamic models from biological data

**DOI:** 10.1101/2023.03.13.532340

**Authors:** Gemma Massonis, Alejandro F. Villaverde, Julio R. Banga

## Abstract

Mechanistic dynamical models allow us to study the behavior of complex biological systems. They can provide an objective and quantitative understanding that would be difficult to achieve through other means. However, the systematic development of these models is a non-trivial exercise and an open problem in computational biology. Currently, many research efforts are focused on model discovery, i.e. automating the development of interpretable models from data. One of the main frameworks is sparse regression, where the sparse identification of nonlinear dynamics (SINDy) algorithm and its variants have enjoyed great success. SINDy-PI is an extension which allows the discovery of rational nonlinear terms, thus enabling the identification of kinetic functions common in biochemical networks, such as Michaelis-Menten. SINDy-PI also pays special attention to the recovery of parsimonious models (Occam’s razor). Here we focus on biological models composed of sets of deterministic nonlinear ordinary differential equations. We present a methodology that, combined with SINDy-PI, allows the automatic discovery of structurally identifiable and observable models which are also mechanistically interpretable. The lack of structural identifiability and observability makes it impossible to uniquely infer parameter and state variables, which can compromise the usefulness of a model by distorting its mechanistic significance and hampering its ability to produce biological insights. We illustrate the performance of our method with six case studies. We find that, despite enforcing sparsity, SINDy-PI sometimes yields models that are unidentifiable. In these cases we show how our method transforms their equations in order to obtain a structurally identifiable and observable model which is also interpretable.

**Author summary:** Dynamical models provide a quantitative understanding of complex biological systems. Since their development is far from trivial, in recent years many research efforts focus on obtaining these models automatically from data. One of the most effective approaches is based on implicit sparse regression. This technique is able to infer biochemical networks with kinetic functions containing rational nonlinear terms. However, as we show here, one limitation is that it may yield models that are unidentifiable. These features may lead to inaccurate mechanistic interpretations and wrong biological insights. To overcome this limitation, we propose an integrated methodology that applies additional procedures in order to ensure that the discovered models are structurally identifiable, observable, and interpretable. We demonstrate our method with six challenging case studies of increasing model complexity.

## Introduction

Mathematical models are increasingly used to describe, monitor, analyze and predict the behavior of complex biological systems. One of the major benefits of using mathematical models to study biology is that they can provide an objective and quantitative understanding that would be difficult to achieve through any other means. In systems biology, dynamical models (typically sets of ordinary differential equations, ODEs) are widely used to provide mechanistic insights into the functioning of biological systems [1, 2].

The use of dynamical systems theory originated in Newtonian mechanics is now pervasive in all the natural and engineering sciences [3]. Dynamic models are highly versatile, enabling researchers to study complex biosystems from a range of different perspectives, such as (i) analyzing the effect of changes in conditions and scenarios different from those studied experimentally, (ii) guiding research by identifying key aspects that need to be further investigated, (iii) helping to generate new testable hypotheses, or (iv) guiding the design of interventions. However, the systematic development of mechanistic dynamic models is a non-trivial exercise. In the case of biological systems, the situation is particularly difficult due to the fact that we cannot rely on first principles in the same way as in e.g. physics. As a consequence, model development is one of the key open problems in mathematical biology [4].

Can we automate the development of mechanistic models? This question of model discovery (in the sense of symbolic reconstruction of equations) from data was already addressed by pioneering attempts in the field of artificial intelligence several decades ago [5–7]. However, the data-driven automatic identification of nonlinear dynamic models has only been addressed more recently. In this area, several different statistical and machine learning frameworks have been considered, including symbolic regression [8, 9], grammar-based methods [10, 11], sparse regression [12], neural networks [13–15], Gaussian process regression [16, 17] and Bayesian approaches [18–20]. More detailed reviews can be found in [21–25]. The sparse identification of nonlinear dynamics (SINDy) algorithm [12] has been particularly successful, and a number of extensions have been developed (see review in [26]),

In the case of biological systems, a large amount of research has been devoted to different classes of subproblems with different simplifying assumptions (such as e.g. static networks, non-mechanistic dynamic networks, linear dynamics, etc.), as reviewed by [27–29]. In this work, we consider the more general problem of fully reconstructing interpretable (mechanistic and parameterized) nonlinear dynamic models from time-series data. Recently, several approaches using methods based on sparse regression, Bayesian identification or symbolic regression have appeared [18, 30–36]. In this context, SINDy-PI [37] is an especially interesting parallel implicit version of SINDy because it allows the incorporation of implicit dynamics and rational nonlinear terms, thus enabling the discovery of kinetic functions (such as Michaelis-Menten) common in biochemical networks.

Many of these SINDy-based methods pay special attention to the recovery of parsimonious models, usually penalizing model complexity [38] or evaluating performance on a validation data-set [37]. The objective is to find the simplest model which can explain the data, in agreement with the well known principle of Occam’s razor. These strategies help to discard more complex models which would be indistinguishable (i.e. would explain the data equally well but adding spurious terms). Besides enforcing simplicity, a related key aspect in model discovery is ensuring structural identifiability and observability (SIO). The property of structural identifiability refers to the theoretical possibility of inferring the unknown parameters of a given model (assuming that its equations are known, except for the numerical values of the parameters) from observations of the model output, which typically consist of time-resolved measurements of its state variables, or of a subset of them [39]. Likewise, observability is the possibility of inferring all the state variables of a model at a given time from future observations of a subset of them. Since lack of SIO makes it impossible to uniquely infer parameters and state variables, it can compromise the usefulness of the model [40–46]. The analysis of these properties can be performed with symbolic computation tools [47], and numerical approaches have also been proposed for their study [48, 49]. However, to the best of our knowledge, ensuring SIO has not been considered in dynamic model discovery yet.

Here we present a methodology that ensures SIO in automatic model discovery in two possible scenarios: with and without prior knowledge. In both cases the end product is a dynamic model of a biological system consisting of (typically nonlinear) ODEs. The equations may contain rational terms, such as Michaelis-Menten kinetics, thus being suitable for the description of many biochemical processes. If there is no prior knowledge about the model structure, the methodology performs equation discovery with the SINDy-PI approach, and incorporates a SIO analysis as a post-processing stage. If there is prior knowledge (i.e. we have a candidate model), another SIO analysis is added as a pre-processing step. If the analyses reveal structural unidentifiabilities, a reparameterization step is carried out to ensure that the resulting model is fully identifiable and observable. Furthermore, equivalent model reformulations are generated to facilitate its interpretation in a mechanistic sense.

Using representative case studies, we illustrate how ignoring these structural properties can lead to wrong conclusions or poorly identified models. Although we demonstrate the use of the methodology with SINDy-PI, it is straightforward to apply it in combination with other automatic discovery methods. In particular, it could easily be adapted to future methods capable of considering partially-observed systems.

## Methods

In this section we describe the methodology, which can be used in two different scenarios. Both of them entail performing model discovery (using SINDy-PI or a similar approach) and performing SIO analysis. If a model is structurally identifiable and observable, we say that it is FISPO (full input, state, and parameter observability). If the SIO analysis reveals that the model is not FISPO, our method suggests a reparameterization step. The two scenarios and their procedures are as follows:

- Scenario (I): full model discovery from time-series data with no prior knowledge. Since we assume zero prior knowledge, we use SINDy-PI to discover a candidate model (CM). We then analyse its SIO. If it is not FISPO, we reparameterize it in order to obtain an equivalent model which is FISPO. Finally, we check if the model is *interpretable*, in the sense that it contains monomials and simple rational terms which belong to a dictionary of mechanistic kinetic terms. If not, we apply a symbolic reformulation step in order to render it interpretable.
- Scenario (II): model discovery from time-series data with prior knowledge. This scenario corresponds to situations where we seek model (in)validation and/or refinement. We assume good prior knowledge and time-series data, that is we are reasonably confident that our prior model (PM), which already represents the data quite well, is close to the ‘true’ one. Here the motivation to use SINDy-PI is to compare this PM with an alternative candidate obtained via model discovery (CM). To this end we check the SIO of the PM, obtaining a reparameterized version if needed. In parallel, we apply SINDy-PI to the data, obtaining a CM, and we make sure that it is FISPO (using reparameterisation if not). If needed, we use model reformulation techniques to obtain interpretable versions of the CM and the PM. Finally, we perform a comparative analysis of these latter models.

An schematic diagram of our method considering these two scenarios is depicted in Fig. (1). In the remainder of this section we describe in detail each of the steps.

**Fig 1.**
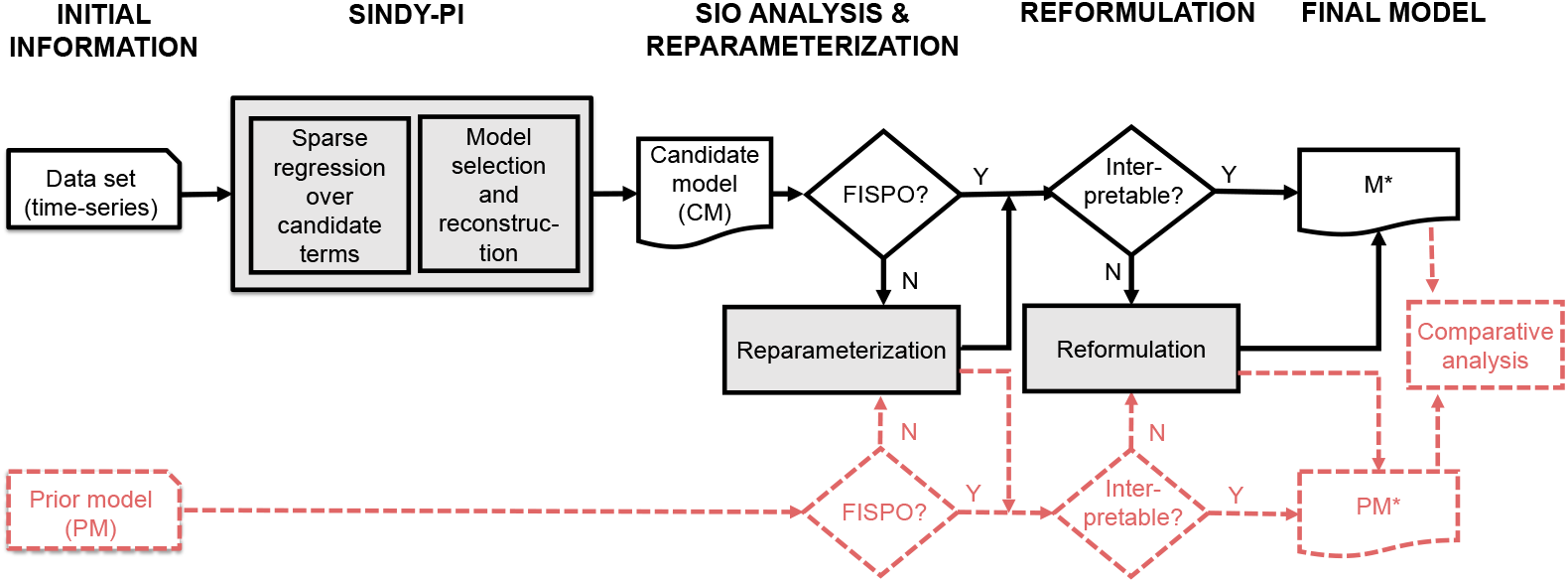
Workflow of the methodology. <u>Scenario (I)</u> (**solid black lines only**): data-driven full model discovery from (time-series) data with no prior knowledge. We apply SINDy-PI and test the SIO of the discovered candidate model (CM). If the CM is not FISPO, we reparameterize it. Next, we check if the model is interpretable; if not, we reformulate it via symbolic manipulation. The result is a FISPO interpretable model, M*. <u>Scenario (II)</u> (solid black lines + dashed, dark orange lines in the lower part): model discovery from (time-series) data with good prior knowledge. In this scenario we seek model (in)validation and/or refinement. We have a prior model (PM) which we want to compare with an alternative candidate discovered from data (CM). To this end, we check the SIO of the PM and reparameterize if needed. In parallel, we apply SINDy-PI to the data to obtain a CM, and we make sure it is FISPO (using reparameterisation if not). Then, we use model reformulation techniques to ensure interpretable versions (M* and PM*) if needed. Lastly, we perform a comparative analysis.

### Automatic model discovery using sparse regression

We assume that the dynamical system is governed by classical reaction-rate nonlinear ordinary differential equations with the following form:

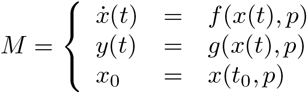

where 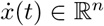 is the state vector, 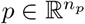 is the parameter vector, the function *f* (*x*(*t*), *p*) represents the dynamics, *y*(*t*) is the measurable output, and *x*_0_ is the vector of initial conditions. SINDy [12] assumes a fully observed system, *y*(*t*) = *x*(*t*). In the reminder of this section, we will consider 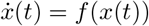 to simplify the notation. SINDy also assumes that *f* (*x*(*t*)) can be expressed as the product of a suitable library function, Θ(*x*(*t*)), and a sparse vector *ξ* (indicating the active library terms), where each entry in the library function is a candidate term:

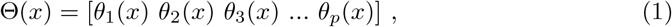

Introducing these new assumptions, 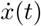 can be expressed as:

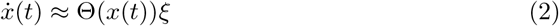

When the system includes rational terms, *f* (*x*) can be rewritten as:

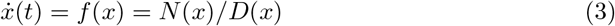

leading to the implicit problem formulation [31]:

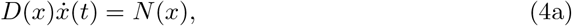

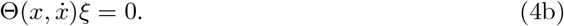

Eq. (4a) has a different kind of term in each side of the equality: the *Left Hand Side (LHS)*, in which there are combinations of term involving the derivative data and the candidate library, and the *Right Hand Side (RHS)*, in which we only have library terms. When *f* (*x*) includes rational terms, model complexity can be viewed as the number of terms in the LHS, as they will involve the denominator degree too.

Once the problem is written in implicit form, Eq. (4b) admits the trivial solution *ξ* = 0. To obtain a solution *ξ ≠* 0 it is necessary to compute the null space, which is an ill-conditioned problem. SINDy-PI [37] is a recent robust extension of SINDy which allows the identification of implicit dynamics and rational nonlinear terms. This variant includes a robust algorithm resulting from assuming that we know at least one term of the library. Then, Eq. (4b) can be expressed in a non-implicit form [37]:

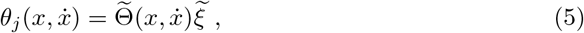

denoting with∼the matrix with the j^*th*^ entry removed, and with 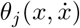 the removed term of the library of functions. To find the sparsest vector *ξ* we solve the following optimization problem:

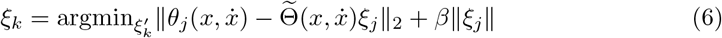

where *β* is the sparsity promoting parameter [12]. One way to relax this non-convex optimization problem is to consider a sequentially thresholded least-squares in which, at each iteration, all the entries below a certain hyperparameter *λ* are set to zero. This threshold must be tuned for each model and each state as it provides different models for the same data set. The optimization process tests each candidate function using brute-force search, which results in a model for each *θ*_*j*_. At the end of this process, we have different models depending on the threshold and the candidate function.

In order to identify the model that best supports the data, SINDy and its variants have explored different model selection methods. Here, we have followed the approach in [31] where a Pareto front is used to find the best trade-off between model complexity and accuracy.

### Structural identifiability and observability analysis and reparameterization

Once a candidate model structure (CM) has been discovered, the next step is to analyse its structural identifiability and observability (SIO) [50]. This test assesses the possibility of determining the values of the model parameters and state variables, respectively, from output measurements. These properties are *structural* (i.e. they depend only on the model equations) and hence they can be analysed *a priori* (i.e. before taking experimental measurements) using symbolic computation. They should not be confused with the so-called *practical* versions of these properties, which depend on the features of the experimental data and are analysed *a posteriori*, i.e. after performing measurements [39].

We can provide a mathematical definition of *structural local identifiability* (SLI) as follows. Let us denote by *y*(*t, p*) the output vector obtained with a parameter vector *p* at time *t*. (For fully observed systems *y*(*t, p*) = *x*(*t, p*), while for partially observed systems *y* typically consists of a subset of *x*.) We say that a parameter *p*_*i*_ (which is the *i*^th^ element of the parameter vector 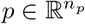 is structurally locally identifiable (SLI) if, for almost any parameter vector 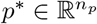, there is a neighbourhood 𝒩 (*p*^*∗*^) such that:

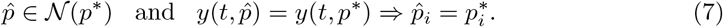

The definition of structural *global* identifiability is similar, but with the neighbourhood 𝒩 (*p*^*∗*^) extending to the whole parameter space. In this paper we focus on SLI.

There are several approaches for determining structural local identifiability and observability. We apply a differential geometry approach, which we explain briefly in these paragraphs. In this framework, parameters are treated as state variables that happen to be constant, i.e. the state vector is augmented with the parameters, 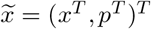, and has dimension 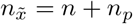. The augmented dynamic equations are 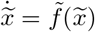, and the output function is 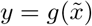, omitting the dependence on time for ease of notation.

Thus, SLI is considered as a particular case of a more general property, observability, which describes the possibility of inferring the internal state of a model by observing its output vector—hence the use of the term FISPO for “full input, state, and parameter observability” (note that this concept also allows for the treatment of unknown inputs as additional state variables, a possibility that we will not consider in this paper).

We analyse SIO by building an observability-identifiability matrix and computing its rank. The matrix is built with Lie derivatives of the output function. The zero-order Lie derivative is 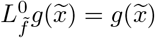, and for *i* ≥ 1 the *i−*order Lie derivatives are obtained as:

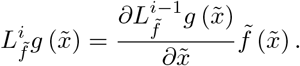

The observability-identifiability matrix 𝒪_*I*_ is:

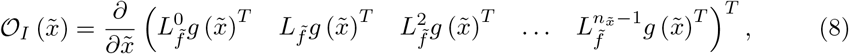

A model is FISPO around a point 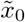 if the rank of its observability-identifiability matrix equals the number of its states and parameters, rank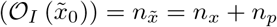. If the rank is smaller, the model contains structurally unidentifiable parameters. By performing additional tests it is possible to determine which specific parameters are structurally identifiable, and which state variables are observable.

If a model is not FISPO, its calibration will almost surely produce wrong parameter estimates. Furthermore, structural unidentifiability is often linked with non-observability, in which case the simulations of some state variables will also be wrong. Thus, structural non-identifiability and non-observability are undesirable features of a model’s structure, which compromise its reliability as a source of biological insight. These features are caused by symmetries in the differential equations of the model that make its output invariant with respect to certain changes in their parameters and/or state variables [51–53]. Said symmetries can be studied in the framework of Lie group theory. We say that a mapping of the form

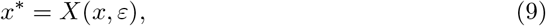

is a one-parameter Lie group of transformations (with *ε* being the parameter) if it has the following properties: it is smooth in *x* and analytic in *ε*, it satisfies the four group axioms (closure, associativity, and existence of an identity and an inverse), and the identity element can be chosen as *ε* = 0. The transformation above is also called a *symmetry transformation*, or a Lie symmetry. Examples of the simplest and possibly most common symmetries in biological modelling include the following:

Translation:

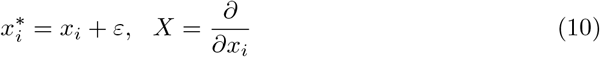

Scaling:

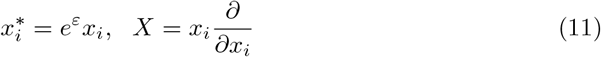

Moebius:

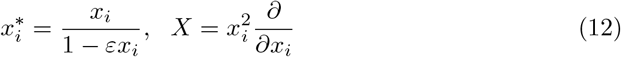

It is sometimes possible to remove or ‘break’ these symmetries by transforming the model equations via a suitable reparameterization. To this end, we first search for the symmetry transformations admitted by the model. If a model has such symmetries, it is overparameterized and therefore structurally unidentifiable. Then, we express the *ε* of those transformations in terms of other parameters, thus setting the value of one of the transformed parameters to one and removing it from the equations. The end result of the reparameterization is a FISPO model that has exactly the same dynamic behaviour as the original one. In previous work [54] we presented a methodology to perform such reparameterizations automatically, which has been integrated in the workflow described here.

In summary, if the SIO analysis of the CM reveals structural unidentifiability and/or non-observability, our methodology applies a symmetry-breaking reparameterization that makes it FISPO.

### Model reformulation for interpretability

The dynamic model obtained in the previous step supports the experimental data and is structurally identifiable and observable. However, the rational expressions in 3 may lack a clear biological interpretation. In the case of biological networks, we will need to reformulate expressions of the form *N* (*x*)*/D*(*x*) into terms that belong to a dictionary of interpretable terms.

Our model reformulation procedure seeks to transform it into simple monomials and rational terms that have a mapping with the dictionary of kinetic and regulatory terms compatible with the specific type of biological reaction network under study. Typically, this dictionary will include mass-action kinetics and simple rational functions (e.g. Michaelis-Menten for enzyme kinetics, or Hill for cooperative binding). However, care should be taken in order to ensure that the reformulation does not destroy identifiability and observability. Further, as shown in the case studies below, sometimes these rational terms can have high degrees, complicating model discovery.

Our reformulation procedure makes use of symbolic manipulation and involves the following steps:

1. Obtain the list of *p* non-trivial divisors of the denominator:

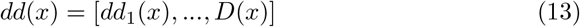
2. Obtain a family A of expressions composed of monomials (interpretable as e.g. mass-action kinetics), minimizing the number of rational terms and their degree, by obtaining the quotients and the residuals:

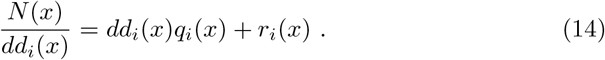
3. If any residuals *r*_*i*_(*x*) lack interpretability, factorize and simplify *N* (*x*) by means of the nested Horner form:

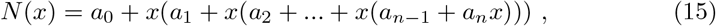

obtaining a family of coupled and factorized equations with the same degree, but with monomials involving different state combinations:

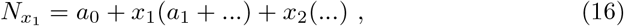

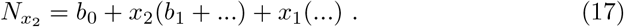

Thus, the Horner nested form gives different possible decompositions of the numerator. Next, obtain a family B of reformulations by simplifying the rational terms using the divisors of Eq. (13) and the Horner form of the numerator. As an example, consider Eq. (18) below, where we can decompose the fraction in the left into a monomial plus a simpler rational term as follows:

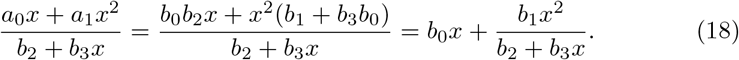
4. Match the monomials and simplified rational terms in families A and B with elements in the dictionary of canonical kinetic and regulatory expressions (or by inspection by a human domain expert), finding members that are fully interpretable.
5. Ensure that the resulting interpretable model is FISPO. If not, reparameterize and repeat until an interpretable and identifiable model is obtained.

### Implementation

We implemented our methodology, as depicted in Fig. (1), using Matlab and the Symbolic Math Toolbox, integrating the following components:

- Sparse regression using SINDy-PI [37] with some modifications, as detailed in the Supporting Information.
- Structural identifiability and observability (SIO) analysis using the algorithm FISPO [55], plus reparameterization using the algorithm AutoRepar [54], as implemented in STRIKE-GOLDD 4.0 and later releases [56].
- Reformulation for interpretability, implementing the algorithm described above using symbolic manipulation.

The resulting code is available at https://doi.org/10.5281/zenodo.7713047. In order to facilitate reproducibility and illustrate the results at each step of the workflow, we have included interactive notebooks (Matlab live scripts) and reports (in HTML format) for each of the case studies described below. More details are given in the Supporting Information.

## Results

Below, we apply our methodology to a set of challenging case studies. In order to illustrate all the steps and capabilities of our workflow, we consider Scenario (II) in all these examples. For each problem, a ground truth (GT) model is defined and subsequently considered as prior model (PM) for the sake of simplicity but without loss of generality. This GT model is used to generate training data sets in all the case studies. After confirming the identifiability and interpretability of the final discovered model M*, we also assess its structural, parametric and predictive accuracy. The predictive power is evaluated taking into account conditions different from those used for generating the training data. Details regarding the training data generation and the conditions to evaluate predictive accuracy are given in Supporting information.

**Table 1.** Main features of the case studies: relevant references and main characteristics of the models considered in the case studies. The fourth and fifth rows show the maximum degree of *N* (*x*) and *D*(*x*) in Eq. (4a). The last row indicates if the original (ground truth, GT) model is fully identifiable and observable (FISPO).

| ID (short name) | Lorenz | Immunity | Bacterial | Microbial | Crypt | Glycolysis |
| --- | --- | --- | --- | --- | --- | --- |
| References | [12,57] | [58] | [31,59] | [60] | [61] | [31,37,62] |
| # states | 3 | 2 | 2 | 2 | 3 | 7 |
| # parameters | 3 | 8 | 5 | 4 | 11 | 13 |
| Max degree $N(x)$ | 2 | 3 | 6 | 2 | 3 | 6 |
| Max degree $D(x)$ | 1 | 2 | 6 | 2 | 4 | 4 |
| FISPO | Y | N | Y | Y | N | Y |

### Lorenz system (Lorenz)

This case study involves the well-known Lorenz system [57], which is a classical example of dynamic model with chaotic behaviour. This model was previously used in [12] to demonstrate the original SINDy algorithm. The governing equations describe the dynamics of a fluid layer warmed from below and cooled from above:

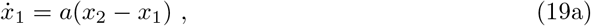

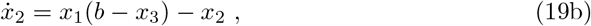

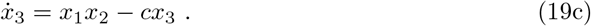

where *x*_1_ is proportional to the rate of convection, *x*_2_ to the horizontal temperature variation and *x*_3_ to the vertical one. For certain values of parameters *a, b*, and *c*, the system exhibits chaotic dynamics.

We consider the ideal Scenario (II) case where the prior model (PM) is the same as the nominal (or ground truth, GT) model. We generate a synthetic training data set using the GT model and settings similar to [12] (details in Supporting Information). Following the workflow in Fig. (1), we perform structural and identifiability analysis and confirm that the PM is fully identifiable and observable (FISPO). We then apply SINDy-PI to the training data, obtaining the following candidate model (CM):

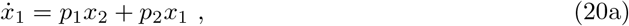

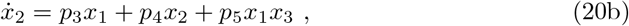

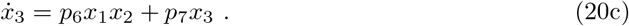

Our algorithm then confirms that this inferred CM model is FISPO and interpretable (thus, it corresponds to M*). Further, it is fully equivalent to the expanded ground truth model in terms of structural, parametric and predictive accuracy, as shown in Fig. (2). In summary, in this case study we have the ideal situation where both the nominal and the inferred models are fully observable and identifiable. As we will see below, this situation might change as soon as we consider rational terms in the dynamics.

**Fig 2.**
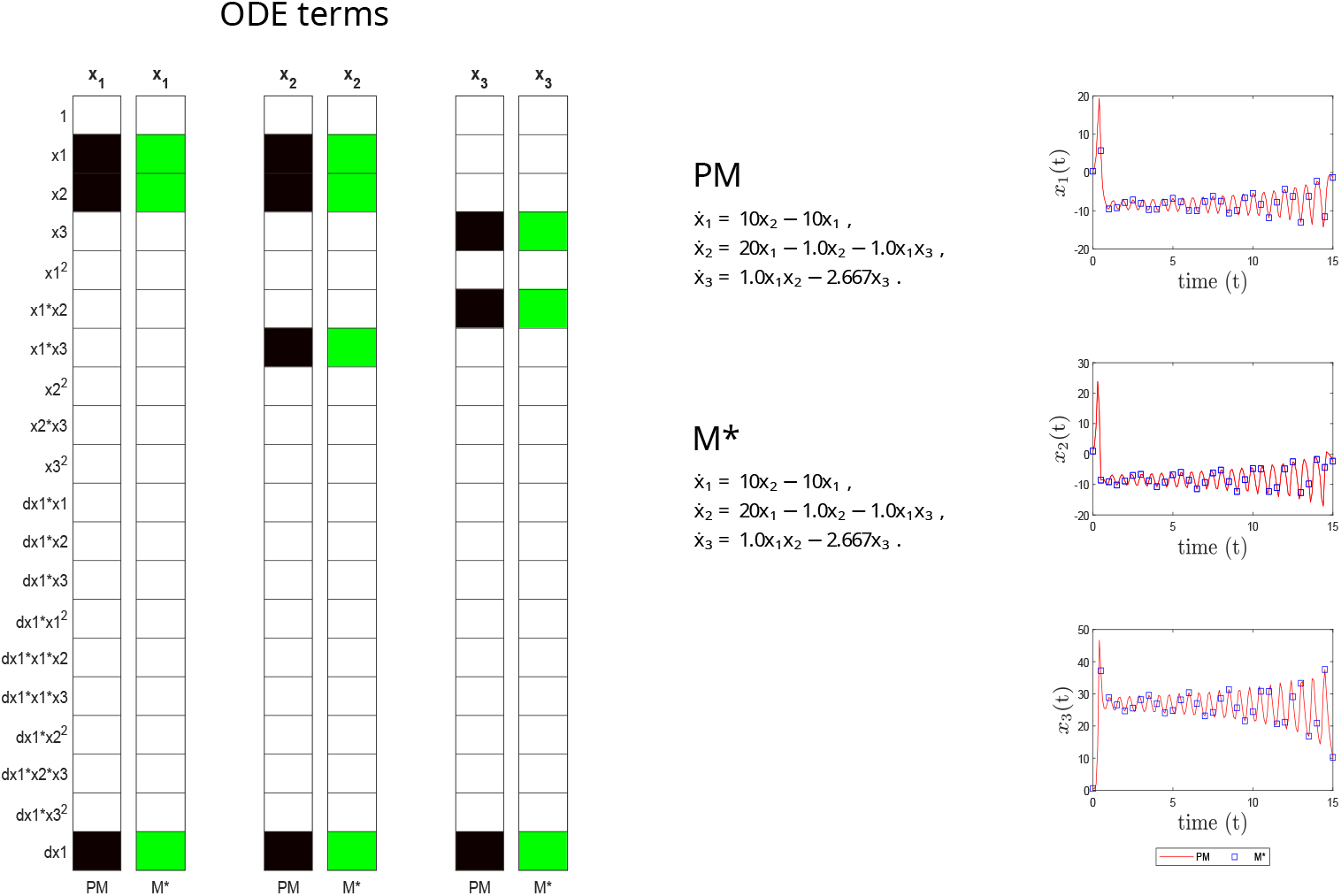
Lorenz case study. Structural accuracy: on the left, active terms in *ξ* (non-zero terms of the prior model PM in black, and of the inferred model M^*^ in green). Parameter accuracy: center, matching parametric ODEs for PM and M^*^. Predictive accuracy: on the right, time evolution of the different states (x_1_, x_2_ and x_3_) of the PM and M^*^ models.

### Competition between bacteria and the immune system (Immunity)

This model describes the influence of quorum sensing signaling molecules (QSSM) on the competition between bacteria and the immune system, as studied in [58]. The following differential equations depict the dynamics for the concentrations of bacteria 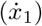 and immune cells 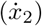:

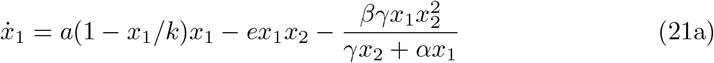

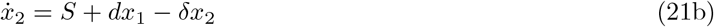

where it is assumed that bacteria grow logistically at rate *a* with an effective carrying capacity of the environment given by parameter *k*, and that they are cleared by the immune system following a mass action term *ex*_1_*x*_2_. The rational term at the end of Eq. (21a) represents the modulation of QSSM in the competition between bacteria and the immune system. We consider Eqs. (21a–21b) as the GT model.

We consider Scenario (II) again, assuming that the prior model (PM) is the same as the ground truth (GT) or nominal model. Considering that the unknown parameters are *a, k, e, β, γ, α, S, d* and *δ*, the structural identifiability analysis of the PM indicates that two of the three parameters involved in the rational term are unidentifiable. Specifically, there is a scaling symmetry between *γ* and *α*, which is probably the most common type of symmetry in biological models [63]. Our reparameterization step indicates that this issue can be solved by dividing the numerator and denominator by one of the unidentifiable parameters; for example, if we choose *α*, Eq. (21a) will be:

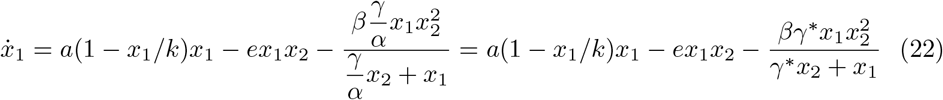

where *γ/α* = *γ*^*∗*^. Thus our new reference model will be the following PM^*^:

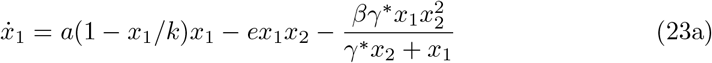

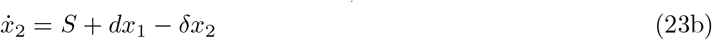

Next, our workflow proceeds by applying SINDy-PI to a data set generated with the GT model, obtaining the following candidate model (CM):

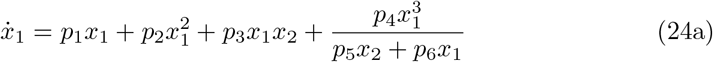

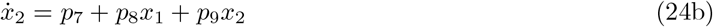

Interestingly, our method then finds that this CM is not FISPO due to three structurally unidentifiable parameters: *p*_4_, *p*_5_, *p*_6_. The reformulation step is then able to find structurally identifiable reformulations of the form:

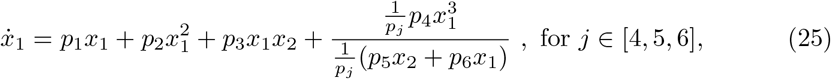

We chose *j* = 6, but the same result can be obtained with *j* = 4, 5. Denoting as 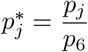, *j* = 4, 5, the resulting dynamic system becomes identifiable. Eq. (25) is re-arranged as:

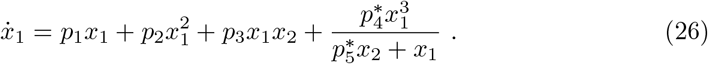

The resulting model is fully identifiable, but the rational term does not match the one in the GT describing the modulation of QSSM. However, the reformulation step in our workflow produces an interpretable form M^*^ which is FISPO:

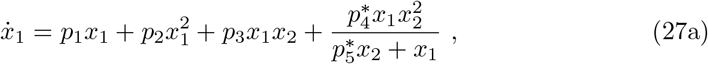

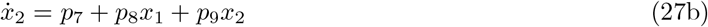

Fig. (3) illustrates the importance of ensuring structural identifiability and observability. The CM found by SINDy-PI is not FISPO, so there are other different parameter realizations producing exactly the same output, as shown by CM2. This means that if this CM structure is used for parameter identification, the estimated parameters will not be unique, i.e. there exist different parameterizations of CM in full agreement with the same output measurements. However, the reformulated model M^*^ is FISPO, i.e. there is a unique set of parameter values compatible with the output. Finally, in Fig. (4) we confirm the structural, parametric and predictive accuracy of M^*^.

**Fig 3.**
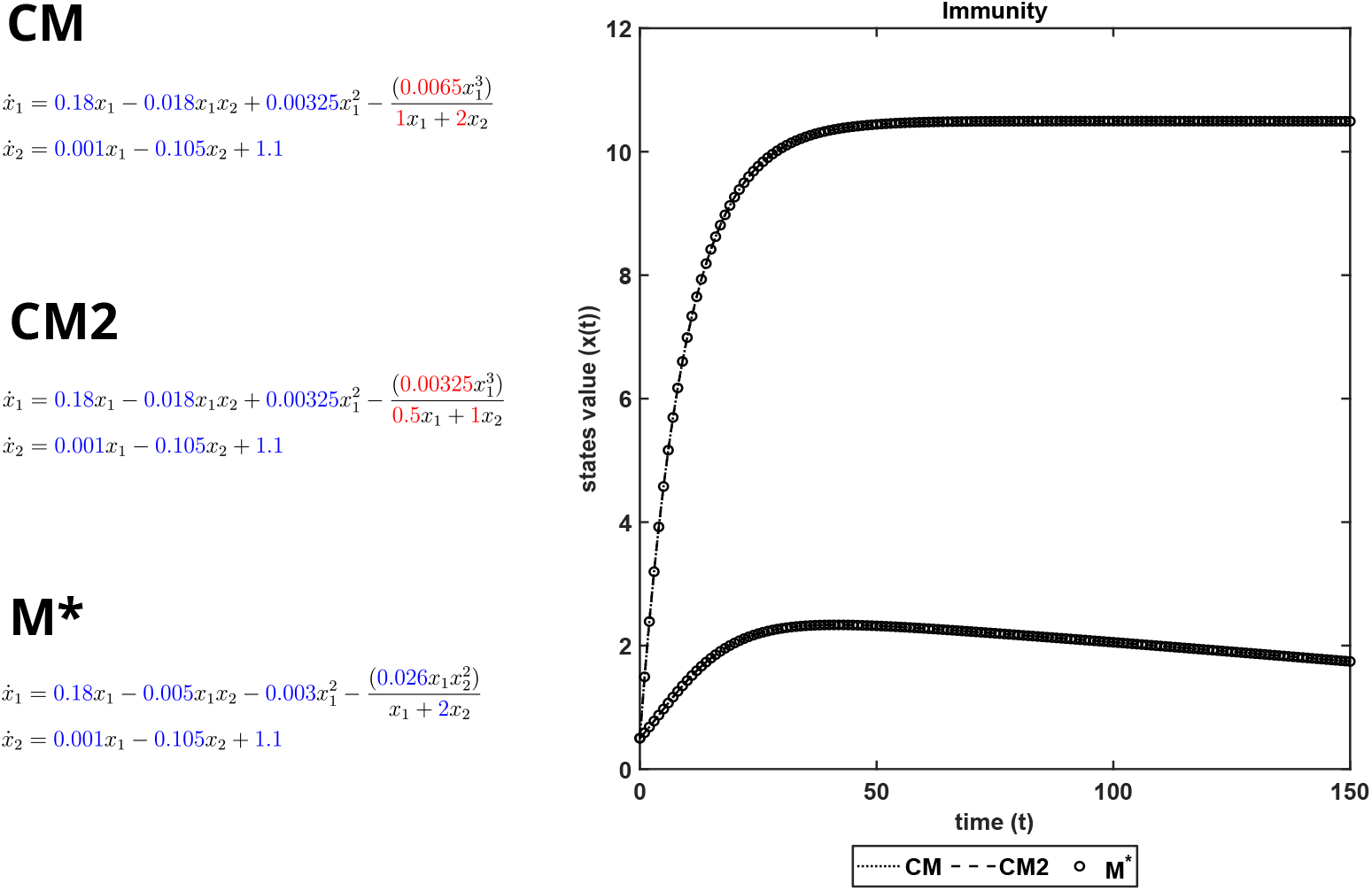
Immunity model. Structural unidentifiability in CM (unidentifiable parameters in red, identifiable parameters in blue) leads to the same output dynamics when different parameterizations are considered, as can be seen in CM2. In contrast, the reformulation M* is FISPO and therefore there is a unique set of parameters compatible with the output measurements.

**Fig 4.**
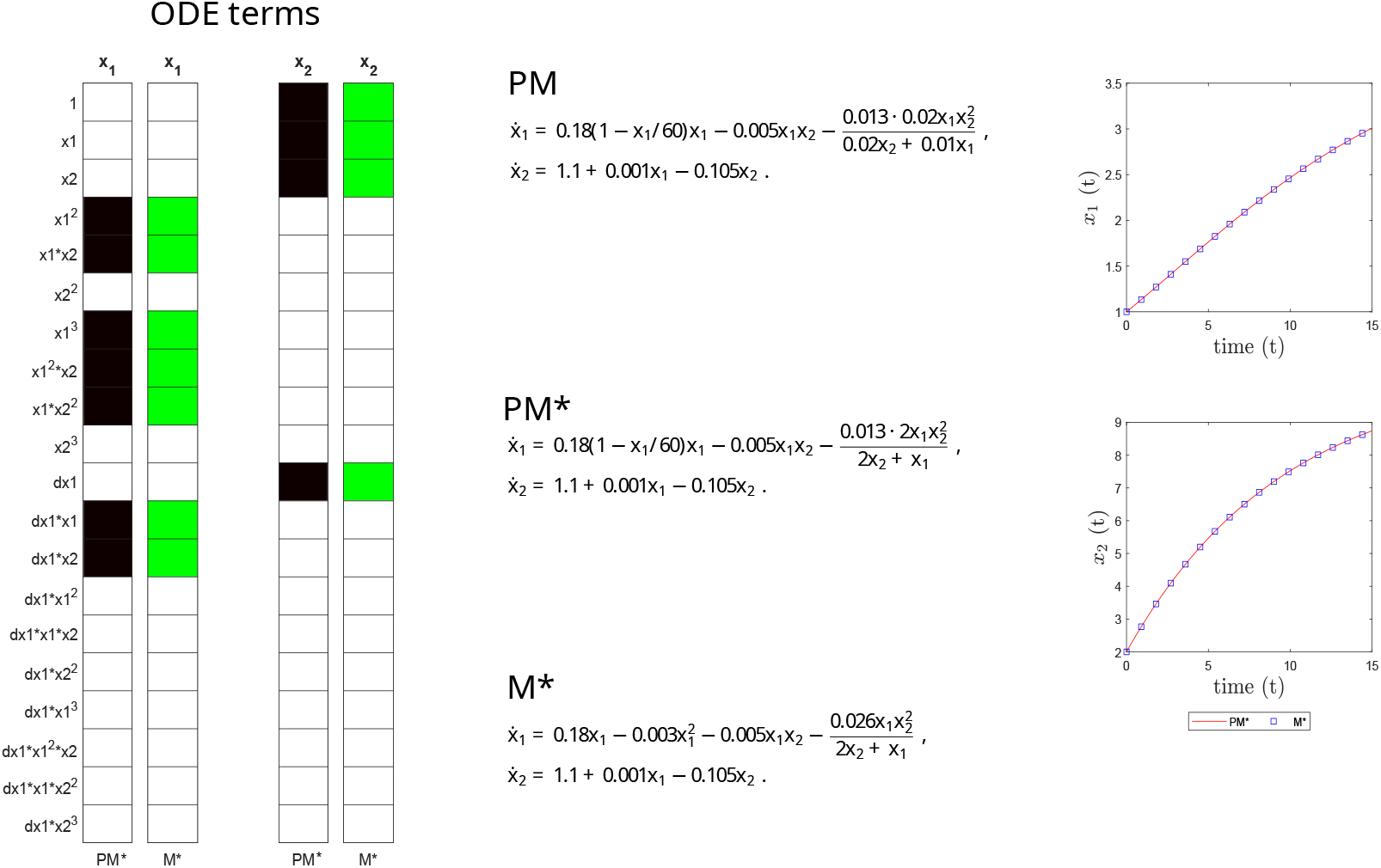
Immunity case study. Structural accuracy: on the left, active terms in *ξ* (non-zero terms of the prior model PM in black, and of the inferred model M^*^ in green). Parameter accuracy: center, matching parametric ODEs for PM and M^*^. Predictive accuracy: on the right, time evolution of the different states (x_1_ and x_2_) of the PM and M^*^ models.

### Stress response in bacteria (Bacterial)

This model describes the stress response in *Bacillus subtilis* [59]. *It was used by Mangan et al [31] to illustrate how an implicit SINDy approach was able to infer biological nonlinear dynamics. Under nutrient limitation, the majority of B. subtilis* cells switch to sporulation, but a small fraction switch to an alternative behaviour, the so called state of competence, in which they are capable of taking up extracellular DNA. This latter fraction might subsequently return to vegetative growth. Süel et al [59] described the regulatory system of this mechanism using a dynamic model with two states. In dimensionless form, the ground truth (GT) model for this example is:

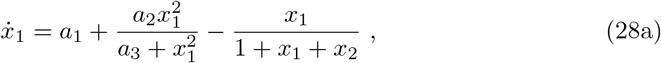

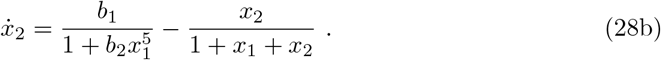

where *x*_1_ and *x*_2_ represent the concentration levels of the ComK and ComS proteins. The rational terms arise from time-scale separation assumptions about the regulation: an autoregulatory positive feedback loop of ComK plus and indirect negative feedback loop mediated by ComS. In Eq. (28a), *a*_1_ corresponds to the minimal rate of ComK production. The second term describes the autoregulation (via a positive feedback loop) of ComK activating its own production, where *a*_2_ is the fully activated rate of ComK generation. The first term in Eq. (28b) describes the negative feedback loop regulating the repression of ComS, where *b*_1_ is the maximum rate of ComS expression. Both the auto-activation of ComK and the repression of ComS follow Hill kinetics where the exponent indicates the level of cooperativity (2 and 5, respectively). The last term in both equations represents the degradation of both ComK and ComS.

We again consider PM=GT and check the identifiability of PM. Considering unknown parameters *a*_1_, *a*_2_, *a*_3_, *b*_1_ and *b*_2_, the model is fully identifiable, thus PM*=GT.

Next, SINDy-PI is applied to the training data generated using GT, obtaining the following candidate model (CM):

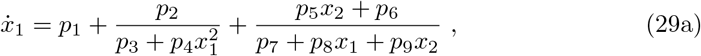

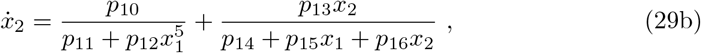

This CM has 16 parameters, *p*_*j*_, *j* = 1, …, 16, and the SIO analysis reveals that all of them are non-identifiable, with the exception of *p*_1_. The reformulation step indicated that we can obtain an identifiable model with four scaling transformations, one per rational term, i.e. the second term in Eq. (29a) is scaled by *p*_*j*_, *j ∈* [2, 3, 4], and the third term by *p*_*k*_, *k ∈* [5, …, 9]. In Eq. (29b) the same strategy is applied for *p*_*l*_, *l ∈* [10, 11, 12], and *p*_*m*_, *m ∈* [13, …, 16]. That is:

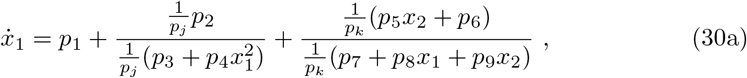

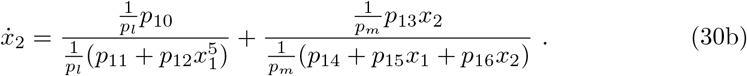

Choosing *j* = 4, *k* = 7, *l* = 11, *m* = 14, the resulting structurally identifiable model is:

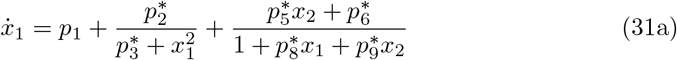

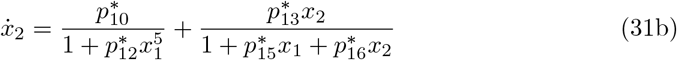

where ^*∗*^ denotes a reparameterized parameter.

This reformulated model is now fully identifiable, but no longer directly interpretable: Eq. (31a) does not explicitly have the term involving the autoregulation of ComK. By means of the reformulation procedure, we are able to recover the autoregulation and degradation terms as in Eq. (28a):

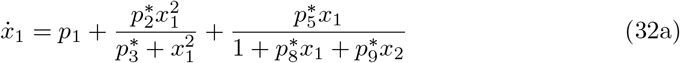

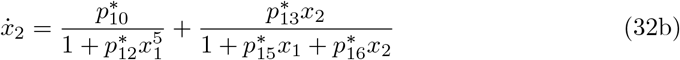

This reformulated model M* is structurally identifiable and interpretable, and equivalent to the PM. Fig. (5) shows the assessment of the structural, parametric and predictive accuracy of the inferred model M^*^.

**Fig 5.**
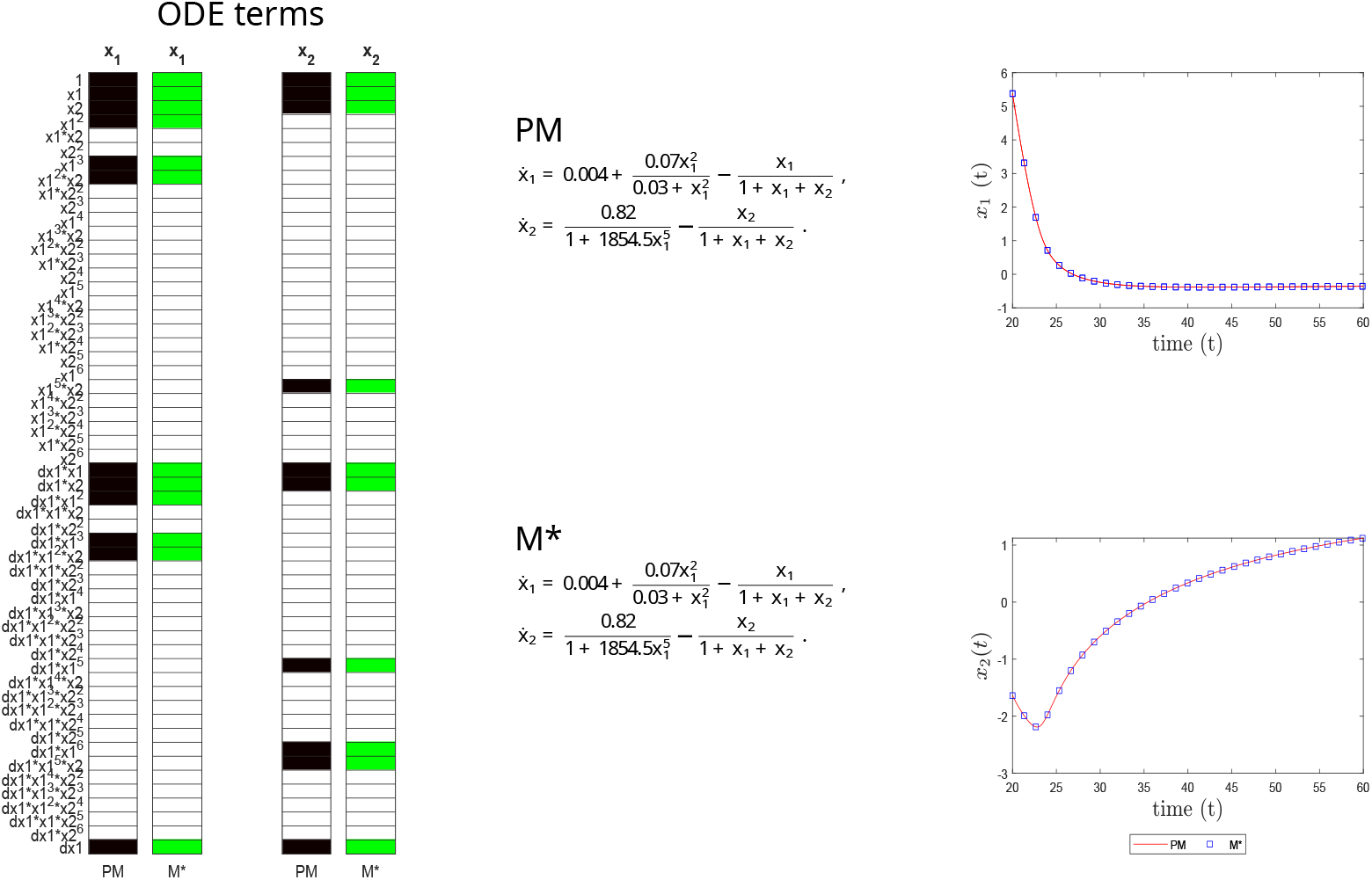
Bacterial case study. Structural accuracy: on the left, active terms in *ξ* (non-zero terms of the prior model PM in black, and of the inferred model M^*^ in green). Parameter accuracy: center, matching parametric ODEs for PM and M^*^. Predictive accuracy: on the right, time evolution of the different states (x_1_ and x_2_) of the PM and M^*^ models.

### Microbial growth (Microbial)

This case study considers microbial growth in a batch reactor, as presented by [64] and later used by [60] to study identifiable reparameterizations of unidentifiable systems. The following model describes the dynamics of microbial and substrate concentrations assuming Monod kinetics (similar in functional form to Michaelis-Menten enzyme kinetics):

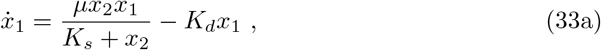

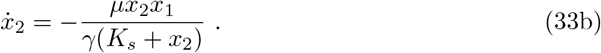

where *x*_1_ and *x*_2_ represent the concentrations of microorganisms and growth-limiting substrate, respectively. The rational term in Eq. (33a) is the Monod kinetic term, where *µ* is the maximum growth velocity and *K*_*s*_ the substrate concentration corresponding to 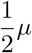. In Eq. (33a), the same rational term appears scaled by *γ* (the yield coefficient) to represent the depletion of substrate. The last term in Eq. (33a) describes the death of microorganisms assuming first order kinetics where *K*_*d*_ is the decay rate.

We consider a prior model (PM) that matches the ground truth (GT) model, Eqs. (33a–33b). When the initial conditions are known and different from zero, our algorithm confirms that this PM is structurally identifiable. Next, our workflow discovers the following dynamics using SINDy-PI:

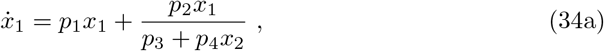

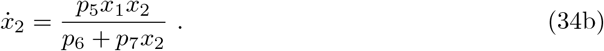

Next, the FISPO step finds that only *p*_1_ is identifiable, i.e. parameters *p*_*i*_, *i* = 2, …, 7 are unidentifiable. The reformulation step finds that it is possible to find an identifiable form by scaling each rational term by the same unidentifiable parameter. For simplicity, we have chosen that 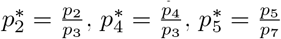 and 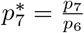. Then, the resulting structurally identifiable model is:

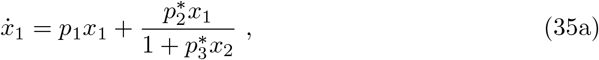

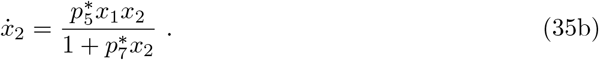

However, Eqs. (35a–35b) are not directly interpretable because they do not contain the expected Monod kinetics terms explicitly. Next, the reformulation step finds an equivalent structure which is both interpretable and identifiable (M*):

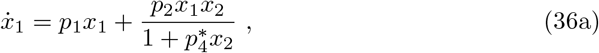

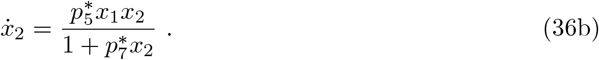

This inferred model (M^*^) is compared to the ground truth in Fig. (6), confirming its structural, parametric and predictive accuracy.

**Fig 6.**
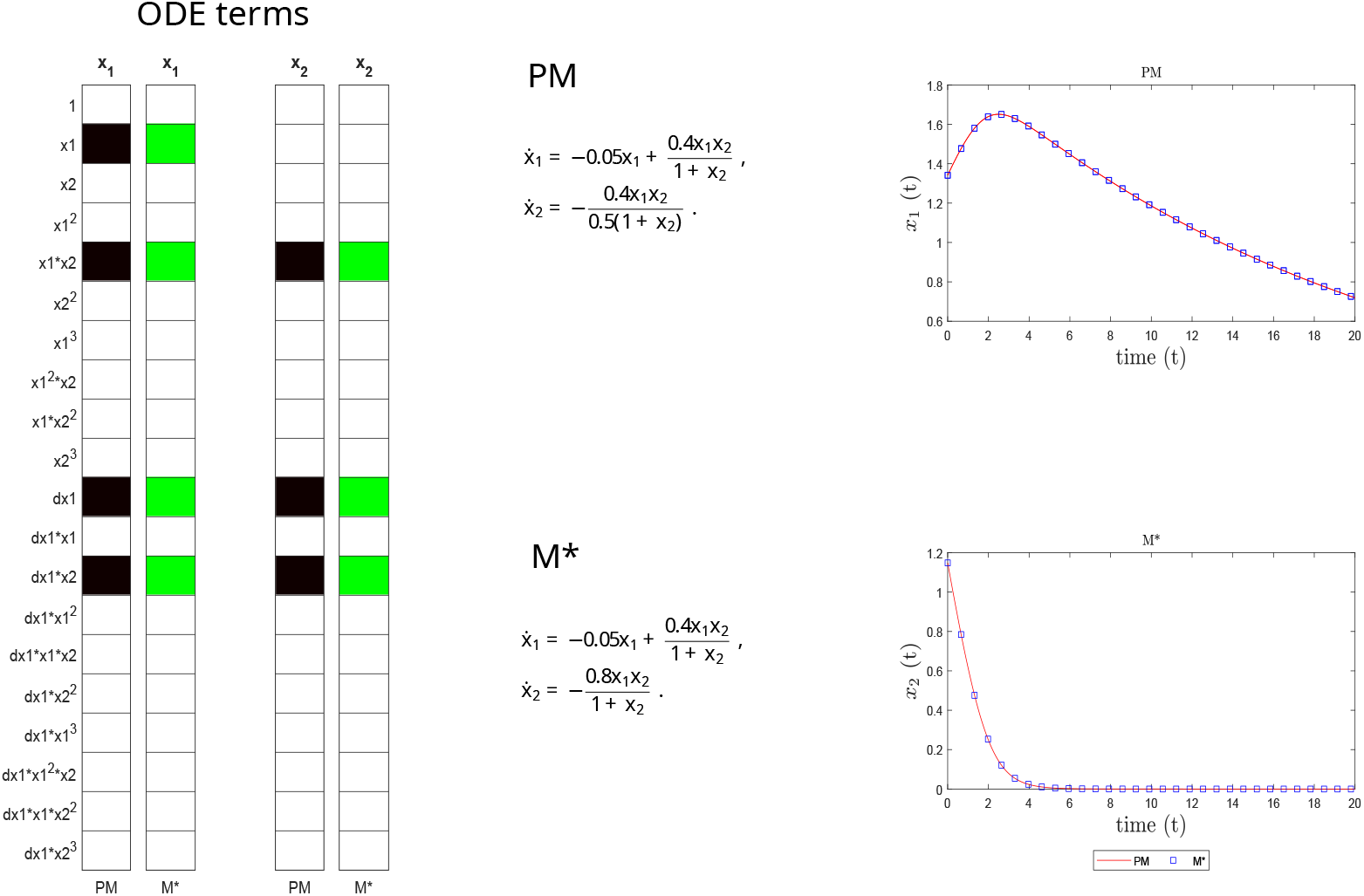
Microbial case study. Structural accuracy: on the left, active terms in *ξ* (non-zero terms of the prior model PM in black, and of the inferred model M^*^ in green). Parameter accuracy: center, matching parametric ODEs for PM and M^*^. On the right, predictive accuracy: time evolution of the different states (x_1_ and x_2_) of the PM and M^*^ models.

### Cell cycle in the colonic crypt (Crypt)

This example considers a cell population model describing the cell renewal cycle in the colonic crypt [61]. This cycle is heavily regulated and the model was used to explain the rupture of homeostasis and the initiation of tumorigenesis. The equations describing the dynamics are:

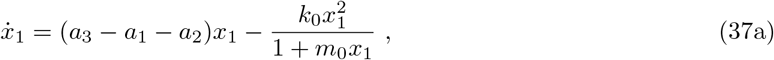

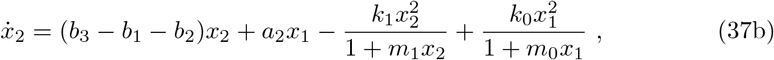

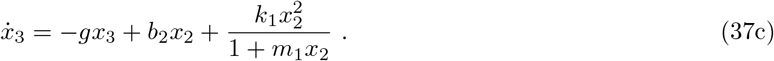

where the state variables represent the populations of stem cells (*x*_1_), semi-differentiated cells (*x*_2_), and fully-differentiated cells (*x*_3_). Stem cells have first order kinetics for renewal (rate given parameter *a*_3_), differentiation (parameter *a*_2_), and death (parameter *a*_1_). Semi-differentiated cells have similar renewal, differentiation and death kinetics (with parameters *b*_*i*_), plus a source term due to the differentiation of stem cells. Fully differentiated cells are generated from semi-differentiated cells with first order rate *b*_2_ and removed with a rate modulated by parameter *g*. The rational terms correspond to saturating feedback mechanism in the differentiation rates.

We take the above model as GT, and PM=GT. Our algorithm finds that this PM is not structurally identifiable: it is not possible to uniquely infer *a*_1_, *a*_3_, *b*_1_ and *b*_3_ due to the presence of a translation symmetry. Next, the reformulation step finds a reparameterized prior model (PM^*^):

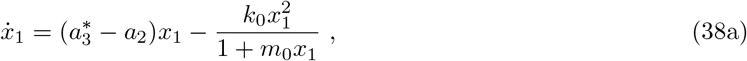

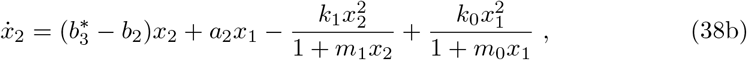

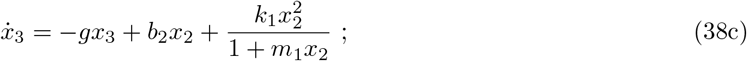

where 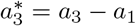 and 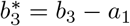. Next, SINDy-PI is applied to the training data, obtaining the following candidate model (CM) :

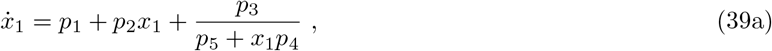

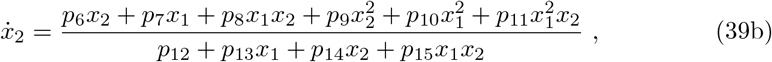

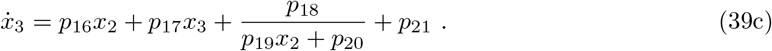

Considering *p*_*i*_, *i* = 1, .., 21 as unknown parameters, the FISPO algorithm indicates that only *p*_1_, *p*_2_, *p*_16_, *p*_17_ and *p*_21_ are structurally identifiable. The reformulation step finds the following structurally identifiable alternative:

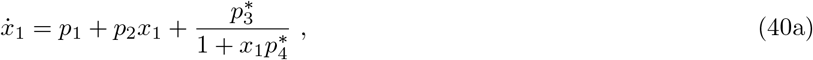

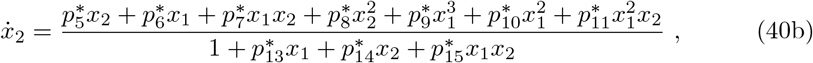

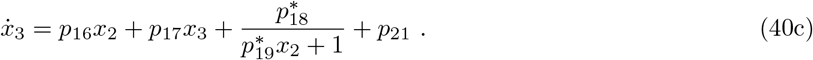

The above model is not directly interpretable, but the reformulation process is able to find the following interpretable and identifiable reformulation M^*^:

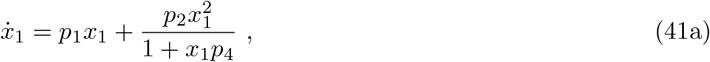

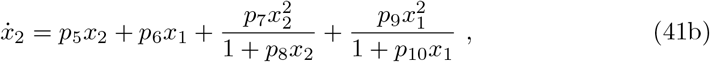

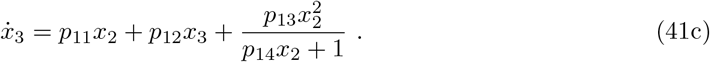

This discovered model M^*^ is fully equivalent to the identifiable version of the ground truth model in terms of structural, parametric and predictive accuracy, as shown in Fig. (7). This example reinforces the importance of checking the identifiability of both the ground truth and the inferred model.

**Fig 7.**
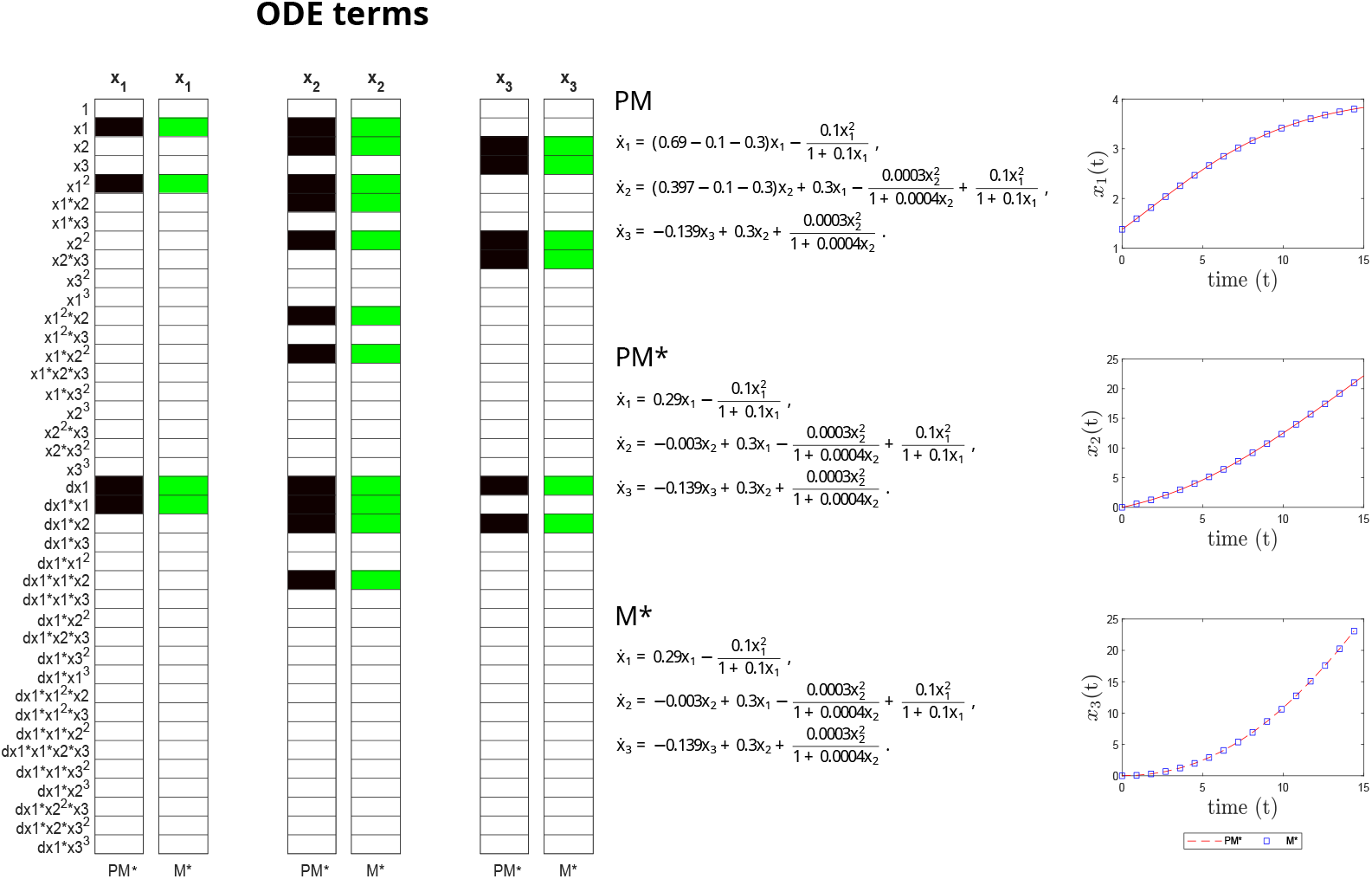
Crypt case study. Structural accuracy: on the left, active terms in *ξ* (non-zero terms of the prior model PM in black, and of the inferred model M^*^ in green). Parameter accuracy: center, matching parametric ODEs for PM and M^*^. Predictive accuracy: on the right, time evolution of the different states (x_1_, x_2_ and x_3_) of the PM and M^*^ models.

### Oscillations in yeast glycolysis (Glycolysis)

Glycolysis is the transformation (in a series of reactions catalyzed by enzymes) of glucose into smaller molecules to produce energy for the cell. In many cell types, glycolysis exhibits oscillations in the concentrations of many intermediate metabolites. This phenomena has been particularly well studied in yeast cells. Wolf and Heinrich [62] studied the oscillatory dynamics of a simplified reaction scheme for yeast glycolysis under anaerobic conditions, where alcoholic fermentation takes place, proposing the following mathematical description:

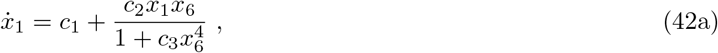

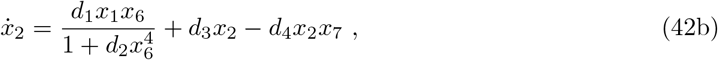

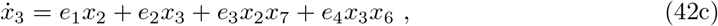

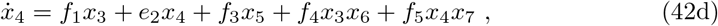

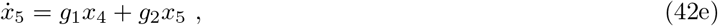

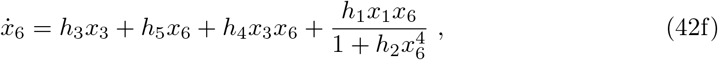

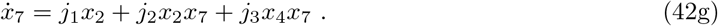

where the state variables represent the concentrations in the cell of glucose (*x*_1_), the pool of triose phosphates (*x*_2_), 1,3-bisphosphoglycerate (*x*_3_), pool of pyruvate and acetaldehyde (*x*_4_), NADH (*x*_5_), ATP (*x*_6_), and *x*_7_ represents the pool of pyruvate and acetaldehyde in the external solution. We consider here the same formulation and parameter values as in [31, 37].

We take the above as GT, and assume PM=GT. Considering all parameters as unknown (*c*_*i*_, *i* = 1, 2, 3; *d*_*i*_, *i* = 1, .., 4; *e*_*i*_, *i* = 1, …, 4; *f*_*i*_, *i* = 1, …, 5; *g*_*i*_, *i* = 1, 2; *h*_*i*_, *i* = 1, …, 5 and *j*_*i*_, *i* = 1, 2, 3), our algorithm confirms that the model is structurally identifiable and observable, i.e. PM*=PM.

This problem is quite challenging for SINDy-PI due to its large number of states and parameters, and the large degree in several terms, leading to a very large library of candidate functions (over 3000 terms). However, it is able to correctly recover the following candidate model (CM):

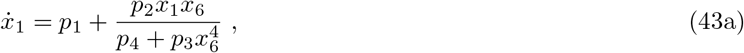

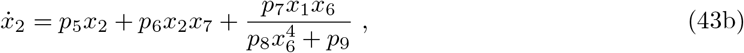

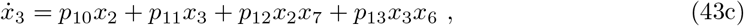

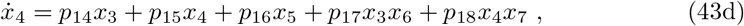

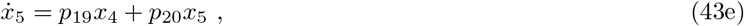

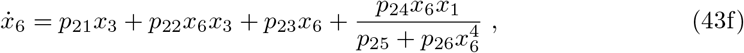

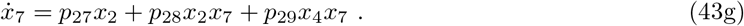

This model has 29 inferred coefficients, *p*_*i*_, *i* = 1, …, 29. Our algorithm analyzes their identifiability and finds that the parameters with indices *i* = 2, 3, 4, 7, 8, 9, 24, 25, 26 are unidentifiable. Next, the reformulation step finds possible reparameterizations by scaling the rational terms. That is, considering the first rational term scaled by *p*_4_, then 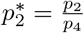 and 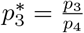; scaling the second term with *p*_9_ produces 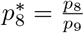 and 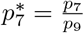 and using *p*_25_ yields 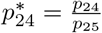 and 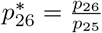 for the last rational term. The end result is an interpretable and identifiable model M*:

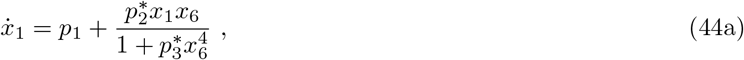

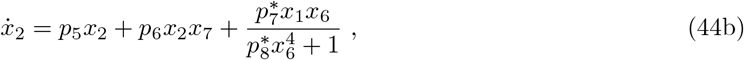

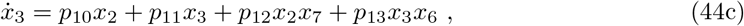

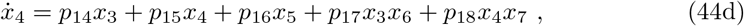

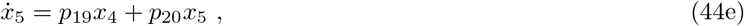

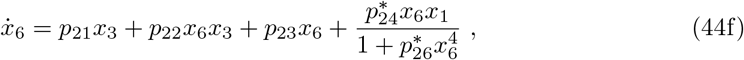

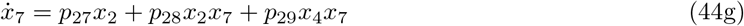

Fig. (8) illustrates the excellent structural, parametric and predictive accuracy of the inferred model.

**Fig 8.**
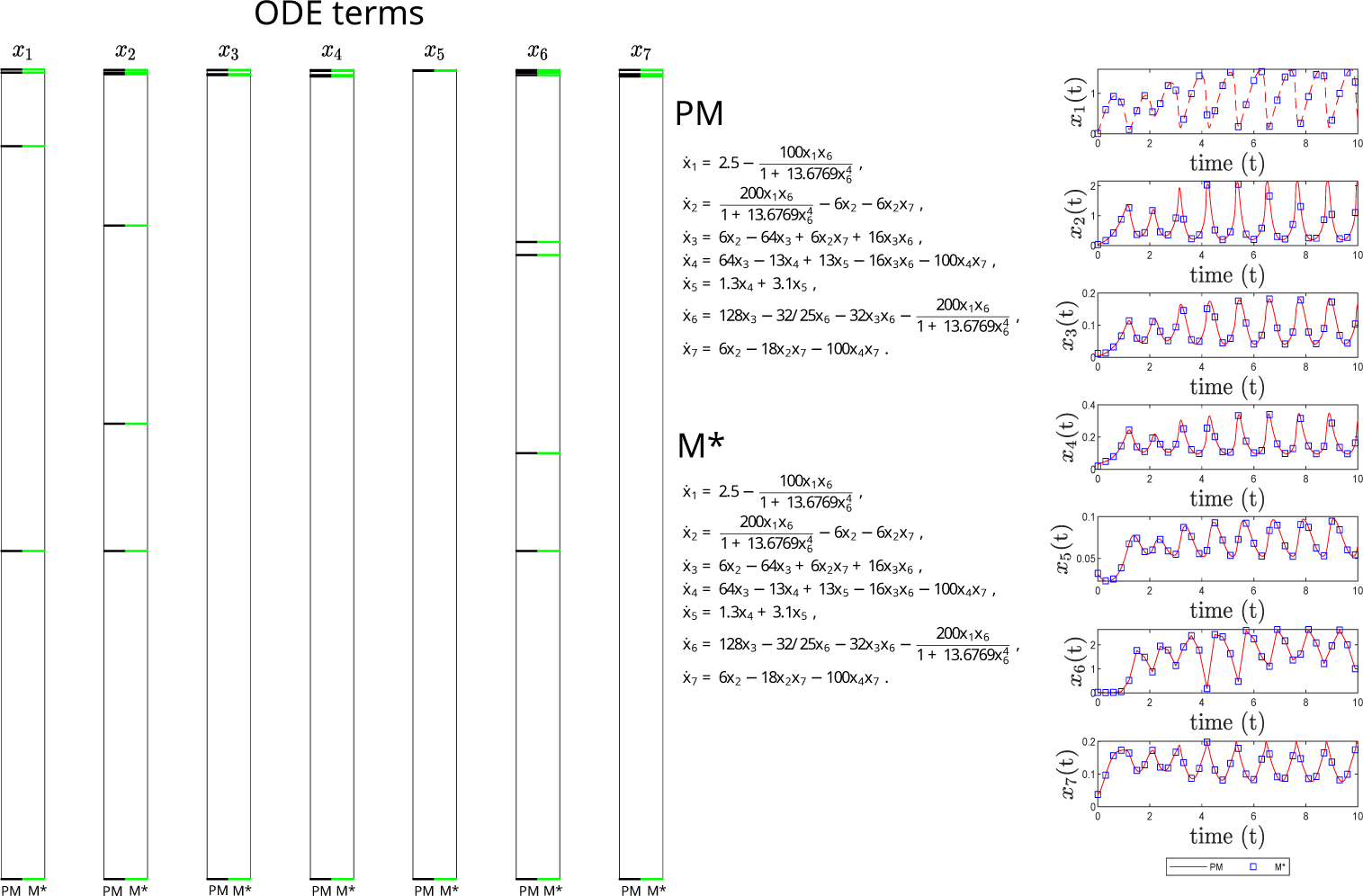
Yeast-Glycolysis case study. Structural accuracy: on the left, active terms in *ξ* (non-zero terms of the prior model PM in black, and of the inferred model M^*^ in green). Due to the large number of terms in *ξ*, the candidate functions are not shown. Parameter accuracy: center, matching parametric ODEs for PM and M^*^. Predictive accuracy: on the right, time evolution of the different states (x_1_, x_2_, x_3_, x_4_, x_5_, x_6_ and x_7_) of the PM and M^*^ models.

## Discussion

In this study we have investigated certain aspects of automatic model discovery techniques to derive mechanistic models of biological systems from time-series data. Specifically, we have focused on possible structural deficiencies of their end result, the inferred model equations. As a reference method we have chosen SINDy-PI [37], a recent sparse regression-based methodology that is particularly suited for computational biology due to its ability to capture complex nonlinearities and rational terms. Since by design SINDy-PI enforces parsimonious models (with the lowest complexity to support the data), it usually produces interpretable equations with excellent predictive power. However, we have shown that sometimes these models lack structural identifiability, which means that using the discovered model structure for parameter estimation might give wrong estimates, compromising its usefulness and reliability.

To address this issue we have presented a methodology that, combined with SINDy-PI, facilitates the inference of identifiable and interpretable dynamic models. Our method integrates symbolic algorithms that analyse a model’s structural identifiability and observability (SIO), reparameterize it to achieve SIO if needed, and reformulate it to make it biologically interpretable. We have illustrated its use in two scenarios, with and without prior knowledge, using six challenging case studies corresponding to different kinds of biological systems, including complex regulatory mechanisms.

Our results highlight additional challenges due to non-obvious issues in the relationship between model reformulation, identifiability and interpretability, and show how our approach is able to successfully surmount them. Importantly, our method is modular and can be easily integrated with other model discovery strategies. While we have demonstrated its application in combination with SINDy-PI, other methods could have been used as well. Furthermore, its calculations are entirely symbolic, i.e. they are not affected by numerical issues caused by insufficient or noisy data (which do however limit the application of the accompanying model discovery method).

Future work will be devoted to model discovery in partially observed systems, where the structural identifiability problem will surely be exacerbated, and observability issues – i.e. the impossibility of inferring some of the unmeasured state variables – are to be expected. It should be noted that, as a matter of fact, our methodology is applicable to partially observed systems in its present form. However, model discovery for such systems is still in its infancy, hence in this study we have considered fully observed systems.

## Data availability

All data and code used for running experiments is available on https://doi.org/10.5281/zenodo.7713047.

## Acknowledgments

This research has received support from grant PID2020-117271RB-C22 (BIODYNAMICS) funded by MCIN/AEI/ 10.13039/501100011033; from the CSIC intramural project grant PIE 202070E062 (MOEBIUS); from grant PID2020-113992RA-I00 funded by MCIN/AEI/ 10.13039/501100011033 (PREDYCTBIO); from grant ED431F 2021/003 funded by Consellería de Cultura, Educación e Ordenación Universitaria, Xunta de Galicia; and from grant RYC-2019-027537-I funded by MCIN/AEI/ 10.13039/501100011033 and by “ESF Investing in your future”. The funding bodies played no role in the design of the study, the collection and analysis of data, or in the writing of the manuscript.

## References

1. DiStefano JJ. Dynamic Systems Biology Modeling and Simulation. Academic Press; 2015.

2. Ingalls BP. Mathematical Modeling in Systems Biology: An Introduction. MIT Press; 2022.

3. Strogatz SH. Nonlinear Dynamics and Chaos: With Applications to Physics, Biology, Chemistry, and Engineering. Westview Press; 2014.

4. Vittadello ST, Stumpf MPH. Open problems in mathematical biology. Math Biosci. 2022;354:108926.

5. Langley P. Data-Driven Discovery of Physical Laws. Cognitive Science. 1981;5(1):31–54.

6. Crutchfield JP, McNamara B. Equations of motion from a data series. Complex systems. 1987;1:417–452.

7. Koza J, Keane MA, Rice JP. Performance improvement of machine learning via automatic discovery of facilitating functions as applied to a problem of symbolic system identification. In: IEEE International Conference on Neural Networks. IEEE; 1993. p. 191–198.

8. Bongard J, Lipson H. Automated reverse engineering of nonlinear dynamical systems. Proc Natl Acad Sci U S A. 2007;104(24):9943–9948.

9. Udrescu SM, Tegmark M. AI Feynman: A physics-inspired method for symbolic regression. Sci Adv. 2020;6(16):eaay2631.

10. Džeroski S, Langley P, Todorovski L. Computational discovery of scientific knowledge. In: Computational discovery of scientific knowledge. Springer; 2007. p. 1–14.

11. Brence J, Todorovski L, Džeroski S. Probabilistic grammars for equation discovery. Knowledge-Based Systems. 2021;224:107077.

12. Brunton SL, Proctor JL, Kutz JN. Discovering governing equations from data by sparse identification of nonlinear dynamical systems. Proc Natl Acad Sci U S A. 2016;113(15):3932–3937.

13. Raissi M, Perdikaris P, Karniadakis GE. Physics informed deep learning (part i): Data-driven solutions of nonlinear partial differential equations. arXiv preprint arXiv:171110561. 2017;.

14. Raissi M, Yazdani A, Karniadakis GE. Hidden fluid mechanics: Learning velocity and pressure fields from flow visualizations. Science. 2020;367(6481):1026–1030.

15. Rackauckas C, Ma Y, Martensen J, Warner C, Zubov K, Supekar R, et al. Universal differential equations for scientific machine learning. arXiv preprint arXiv:200104385. 2020;.

16. Bhouri MA, Perdikaris P. Gaussian processes meet NeuralODEs: a Bayesian framework for learning the dynamics of partially observed systems from scarce and noisy data. Philos Trans A Math Phys Eng Sci. 2022;380(2229):20210201.

17. VandenHeuvel DJ, Drovandi C, Simpson MJ. Computationally efficient mechanism discovery for cell invasion with uncertainty quantification. PLoS Comput Biol. 2022;18(11):e1010599.

18. Pan W, Yuan Y, Gonçalves J, Stan GB. A sparse Bayesian approach to the identification of nonlinear state-space systems. IEEE Transactions on Automatic Control. 2015;61(1):182–187.

19. Zhang S, Lin G. Robust data-driven discovery of governing physical laws with error bars. Proc Math Phys Eng Sci. 2018;474(2217):20180305.

20. Guimerà R, Reichardt I, Aguilar-Mogas A, Massucci FA, Miranda M, Pallarès J, et al. A Bayesian machine scientist to aid in the solution of challenging scientific problems. Sci Adv. 2020;6(5):eaav6971.

21. Džeroski S, Todorovski L. Equation discovery for systems biology: finding the structure and dynamics of biological networks from time course data. Current Opinion in Biotechnology. 2008;19(4):360–368.

22. North JS, Wikle CK, Schliep EM. A Review of Data-Driven Discovery for Dynamic Systems. arXiv preprint arXiv:221010663. 2022;.

23. Willard J, Jia X, Xu S, Steinbach M, Kumar V. Integrating physics-based modeling with machine learning: A survey. arXiv preprint arXiv:200304919. 2020;1(1):1–34.

24. Brunton SL, Kutz JN. Data-driven science and engineering: Machine learning, dynamical systems, and control; 2nd edition. Cambridge University Press; 2022.

25. Ghadami A, Epureanu BI. Data-driven prediction in dynamical systems: recent developments. Philosophical Transactions of the Royal Society A. 2022;380(2229):20210213.

26. Naozuka GT, Rocha HL, Silva RS, Almeida RC. SINDy-SA framework: enhancing nonlinear system identification with sensitivity analysis. Nonlinear Dyn. 2022;110(3):2589–2609.

27. Villaverde AF, Banga JR. Reverse engineering and identification in systems biology: strategies, perspectives and challenges. Journal of the Royal Society Interface. 2014;11(91):20130505.

28. Kirk P, Silk D, Stumpf MP. Reverse engineering under uncertainty. In: Uncertainty in biology. Springer; 2016. p. 15–32.

29. Mercatelli D, Scalambra L, Triboli L, Ray F, Giorgi FM. Gene regulatory network inference resources: A practical overview. Biochimica et Biophysica Acta (BBA)-Gene Regulatory Mechanisms. 2020;1863(6):194430.

30. Sunnåker M, Zamora-Sillero E, Dechant R, Ludwig C, Busetto AG, Wagner A, et al. Automatic generation of predictive dynamic models reveals nuclear phosphorylation as the key Msn2 control mechanism. Science signaling. 2013;6(277):ra41–ra41.

31. Mangan NM, Brunton SL, Proctor JL, Kutz JN. Inferring biological networks by sparse identification of nonlinear dynamics. IEEE Transactions on Molecular, Biological and Multi-Scale Communications. 2016;2(1):52–63.

32. Daniels BC, Ryu WS, Nemenman I. Automated, predictive, and interpretable inference of escape dynamics. Proc Natl Acad Sci U S A. 2019;116(15):7226–7231.

33. Rajewsky N, Jurga S, Barciszewski J. Systems Biology. Springer; 2018.

34. Hoffmann M, Fröhner C, Noé F. Reactive SINDy: Discovering governing reactions from concentration data. J Chem Phys. 2019;150(2):025101.

35. Yeung E, Kim J, Yuan Y, Gonçalves J, Murray RM. Data-driven network models for genetic circuits from time-series data with incomplete measurements. J R Soc Interface. 2021;18(182):20210413.

36. Jiang R, Singh P, Wrede F, Hellander A, Petzold L. Identification of dynamic mass-action biochemical reaction networks using sparse Bayesian methods. PLoS Comput Biol. 2022;18(1):e1009830.

37. Kaheman K, Kutz JN, Brunton SL. SINDy-PI: a robust algorithm for parallel implicit sparse identification of nonlinear dynamics. Proc Math Phys Eng Sci. 2020;476(2242):20200279.

38. Mangan NM, Kutz JN, Brunton SL, Proctor JL. Model selection for dynamical systems via sparse regression and information criteria. Proc Math Phys Eng Sci. 2017;473(2204):20170009.

39. Wieland FG, Hauber AL, Rosenblatt M, T”onsing C, Timmer J. On structural and practical identifiability. Current Opinion in Systems Biology. 2021;25:60–69.

40. Szederkényi G, Banga JR, Alonso AA. Inference of complex biological networks: distinguishability issues and optimization-based solutions. BMC systems biology. 2011;5(1):1–15.

41. Chin SV, Chappell MJ. Structural identifiability and indistinguishability analyses of the Minimal Model and a Euglycemic Hyperinsulinemic Clamp model for glucose–insulin dynamics. Computer Methods and Programs in Biomedicine. 2011;104(2):120–134.

42. Janzén DLI, Bergenholm L, Jirstrand M, Parkinson J, Yates J, Evans ND, et al. Parameter Identifiability of Fundamental Pharmacodynamic Models. Front Physiol. 2016;7:590.

43. Villaverde AF, Banga JR. Dynamical compensation and structural identifiability of biological models: Analysis, implications, and reconciliation. PLoS Comput Biol. 2017;13(11):e1005878.

44. Eisenberg MC, Jain HV. A confidence building exercise in data and identifiability: Modeling cancer chemotherapy as a case study. Journal of theoretical biology. 2017;431:63–78.

45. Muñoz-Tamayo R, Puillet L, Daniel JB, Sauvant D, Martin O, Taghipoor M, et al. To be or not to be an identifiable model. Is this a relevant question in animal science modelling? Animal. 2018;12(4):701–712.

46. Schmidt PJ, Emelko MB, Thompson ME. Recognizing Structural Nonidentifiability: When Experiments Do Not Provide Information About Important Parameters and Misleading Models Can Still Have Great Fit. Risk Anal. 2020;40(2):352–369.

47. Barreiro XR, Villaverde AF. Benchmarking tools for a priori identifiability analysis. Bioinformatics. 2023;39.

48. Raue A, Kreutz C, Maiwald T, Bachmann J, Schilling M, Klingmüller U, et al. Structural and practical identifiability analysis of partially observed dynamical models by exploiting the profile likelihood. Bioinformatics. 2009;25(15):1923–1929.

49. Stigter J, Joubert D. Computing measures of identifiability, observability, and controllability for a dynamic system model with the StrucID App. IFAC-PapersOnLine. 2021;54(7):138–143.

50. Villaverde AF, et al. Observability and structural identifiability of nonlinear biological systems. Complexity. 2019;2019.

51. Yates JW, Evans ND, Chappell MJ. Structural identifiability analysis via symmetries of differential equations. Automatica. 2009;45(11):2585–2591.

52. Merkt B, Timmer J, Kaschek D. Higher-order Lie symmetries in identifiability and predictability analysis of dynamic models. Physical Review E. 2015;92(1):012920.

53. Villaverde AF. Symmetries in Dynamic Models of Biological Systems: Mathematical Foundations and Implications. Symmetry. 2022;14(3):467.

54. Massonis G, Banga JR, Villaverde AF. AutoRepar: a method to obtain identifiable and observable reparameterizations of dynamic models with mechanistic insights. International Journal of Robust and Nonlinear Control. 2021;.

55. Villaverde AF, Tsiantis N, Banga JR. Full observability and estimation of unknown inputs, states and parameters of nonlinear biological models. Journal of the Royal Society Interface. 2019;16(156):20190043.

56. Díaz-Seoane S, Rey Barreiro X, Villaverde AF. STRIKE-GOLDD 4.0: user-friendly, efficient analysis of structural identifiability and observability. Bioinformatics. 2022;39(1). doi:10.1093/bioinformatics/btac748.

57. Lorenz EN. Deterministic Nonperiodic Flow. Journal of the Atmospheric Sciences. 1963;20(2):130–141. doi:10.1175/1520-0469(1963)020<0130:dnf>2.0.co;2.

58. Zhang Z. Mathematical Model of a Bacteria-Immunity System with the Influence of Quorum Sensing Signal Molecule. Journal of Applied Mathematics and Physics. 2016;04(05):888–896. doi:10.4236/jamp.2016.45097.

59. Süel GM, Garcia-Ojalvo J, Liberman LM, Elowitz MB. An excitable gene regulatory circuit induces transient cellular differentiation. Nature. 2006;440(7083):545–550.

60. Evans ND, Chappell MJ. Extensions to a procedure for generating locally identifiable reparameterisations of unidentifiable systems. Mathematical Biosciences. 2000;168(2):137–159. doi:10.1016/s0025-5564(00)00047-x.

61. Johnston MD, Edwards CM, Bodmer WF, Maini PK, Chapman SJ. Examples of Mathematical Modeling: Tales from the Crypt. Cell Cycle. 2007;6(17):2106–2112. doi:10.4161/cc.6.17.4649.

62. Wolf J, Heinrich R. Effect of cellular interaction on glycolytic oscillations in yeast: a theoretical investigation. Biochemical Journal. 2000;345(2):321–334. doi:10.1042/bj3450321.

63. Castro M, de Boer RJ. Testing structural identifiability by a simple scaling method. PLOS Computational Biology. 2020;16(11):e1008248.

64. Holmberg A. On the practical identifiability of microbial growth models incorporating Michaelis-Menten type nonlinearities. Mathematical Biosciences. 1982;62(1):23–43. doi:10.1016/0025-5564(82)90061-x.

