## Supplemental material for "Distilling identifiable and interpretable dynamic models from biological data"

### Supporting Information for "Distilling identifiable and interpretable models from biological data"

Gemma Massonis [1], Alejandro F. Villaverde [2,3] and Julio R. Banga [1]
[1]Computational Biology Lab, MBG-CSIC, Pontevedra, Galicia, Spain,
[2]CITMAga, Santiago de Compostela, Galicia, Spain,
[3]Universidade de Vigo, Dept. of Systems and Control Engineering, Vigo, Galicia, Spain


March 10, 2023

#### 1 Generation of training data sets

The training data sets for the case studies were generated following the guidelines in [1, 2]. A set of $N_{\mathrm{IC}}$ different initial conditions were randomly generated inside a range defined by lower and upper bounds $(lb, ub)$. Data were generated by simulating each model with its nominal parameters and different initial conditions for a certain time horizon $(t_0, t_{\mathrm{f}})$, with sampling (measurement) times given by a time step $dt$. Next, 10% of the total generated data was selected via random permutation as training data. Finally, a different set of initial conditions $(ICS_{pp})$ were subsequently used to evaluate the predictive power of the discovered model. Below, we report the values of these settings for each case study.

##### 1.1 `Lorenz`

| parameter | a | b | c |
|---|---|---|---|
| value | 10 | 28 | 8/3 |

Table 1: `Lorenz`: nominal values of the parameters.

| Setting | value |
|---|---|
| $N_{\mathrm{IC}}$ | 80 |
| $lb$ | 1e-4 |
| $ub$ | 0.4706 |
| $t_0$ | 0 |
| $t_{\mathrm{f}}$ | 15 |
| $dt$ | 0.1 |
| $ICS_{pp}$ | [0.35,0.2,0.4] |

Table 2: `Lorenz`: settings used to generate the training data set and evaluate the predictive power of the discovered model.

#### 1.2 Immunity

| parameter | a | k | e | $\beta$ | $\gamma$ | $\alpha$ | S | d | $\delta$ |
|---|---|---|---|---|---|---|---|---|---|
| value | 0.18 | 60 | 0.005 | 0.013 | 0.02 | 0.01 | 1.1 | 0.001 | 0.105 |

Table 3: `Immunity`: nominal values of the parameters.

| Setting | value |
|---|---|
| $N_{\mathrm{IC}}$ | 80 |
| $lb$ | 0.0500 |
| $ub$ | 75.9614 |
| $t_0$ | 0 |
| $t_{\mathrm{f}}$ | 15 |
| $dt$ | 0.1 |
| $ICS_{pp}$ | [0.2635;0.8278] |

Table 4: `Immunity`: settings used to generate the training data set and evaluate the predictive power of the discovered model.

#### 1.3 Bacterial

| parameter | $a_1$ | $a_2$ | $a_3$ | $b_1$ | $b_2$ |
|---|---|---|---|---|---|
| value | 0.004 | 0.07 | 0.04 | 0.82 | 1854.5 |

Table 5: `Bacterial`: nominal values of the parameters.

| Setting | value |
|---|---|
| $N_{\mathrm{IC}}$ | 80 |
| $lb$ | 1e-4 |
| $ub$ | 4.9 |
| $t_0$ | 0 |
| $t_{\mathrm{f}}$ | 15 |
| $dt$ | 0.1 |
| $ICS_{pp}$ | [0.8;0.9] |

Table 6: `Bacterial`: settings used to generate the training data set and evaluate the predictive power of the discovered model.

#### 1.4 `Microbial`

| parameter | $K_d$ | $K_s$ | $\mu$ | $\gamma$ |
|---|---|---|---|---|
| value | 0.05 | 1 | 0.4 | 0.5 |

Table 7: `Microbial`: nominal values of the parameters.

| Setting | value |
|---|---|
| $N_{IC}$ | 80 |
| $lb$ | 1e-4 |
| $ub$ | 0.4706 |
| $t_0$ | 0 |
| $t_f$ | 15 |
| $dt$ | 0.1 |
| $ICS_{pp}$ | [0.8;0.9] |

Table 8: `Microbial`: settings used to generate the training data set and evaluate the predictive power of the discovered model.

#### 1.5 `Crypt`

| parameter | $a_1$ | $a_2$ | $a_3$ | $b_1$ | $b_2$ | $b_3$ | g | $k_0$ | $k_1$ | $m_0$ | $m_1$ |
|---|---|---|---|---|---|---|---|---|---|---|---|
| value | 0.1 | 0.3 | 0.69 | 0.1 | 0.3 | 0.397 | 0.139 | 0.1 | 3e-4 | 0.1 | 4e-4 |

Table 9: `Crypt`: nominal values of the parameters.

| Setting | value |
|---|---|
| $N_{IC}$ | 80 |
| $lb$ | 1e-4 |
| $ub$ | 79.9582 |
| $t_0$ | 0 |
| $t_f$ | 15 |
| $dt$ | 0.1 |
| $ICS_{pp}$ | [0.35;0.2;0.1] |

Table 10: `Crypt`: settings used to generate the training data set and evaluate the predictive power of the discovered model.

##### 1.6 Glycolysis

| parameter | $c_1$ | $c_2$ | $c_3$ | $d_1$ | $d_2$ | $d_3$ | $d_4$ | $e_1$ | $e_2$ | $e_3$ | $e_4$ | $f_1$ | $f_2$ |
|---|---|---|---|---|---|---|---|---|---|---|---|---|---|
| value | 2.5 | -100 | 13.6769 | 200 | 13.6769 | -6 | -6 | 6 | -64 | 6 | 16 | 64 | -13 |
| **parameter** | $f_3$ | $f_4$ | $f_5$ | $g_1$ | $g_2$ | $h_1$ | $h_2$ | $h_3$ | $h_4$ | $h_5$ | $j_1$ | $j_2$ | $j_3$ |
| value | 13 | -16 | -100 | 1.3 | 3.1 | -200 | 13.6769 | 128 | -32 | -32/25 | 6 | -18 | -100 |

Table 11: `Glycolysis`: nominal values of the parameters.

| Setting | value |
|---|---|
| $N_{\text{IC}}$ | 450 |
| $lb$ | 1e-7 |
| $ub$ | 0.25 |
| $t_0$ | 0 |
| $t_{\text{f}}$ | 15 |
| $dt$ | 0.1 |
| $ICS_{pp}$ | [0.5;0.2;1.1;0.8;0.5;0.4;0.1] |

Table 12: `Glycolysis`: settings used to generate the training data set and evaluate the predictive power of the discovered model.

#### 2 Code availability

The code to reproduce the results of this study is available at `https://doi.org/10.5281/zenodo.7713047`. For each case study, we provide a Matlab interactive notebook (live script) which executes the different steps of our workflow. To facilitate reproducibility, we also provide detailed reports (in HTML format) with the results obtained for each case study.

Requirements:

- MATLAB (tested with version R2020b under Win10), including:
  - Symbolic Math Toolbox.
  - Parallel Computing Toolbox (optional).
- STRIKE-GOLDD (tested with version 4.0.2), available at `https://github.com/afvillaverde/strike-goldd`)
- SINDy-PI, available at `https://github.com/dynamicslab/SINDy-PI.git`.

Installation: please follow the instructions in the `README_install.txt` file.

#### References


[1] Mangan NM, Brunton SL, Proctor JL, Kutz JN. Inferring biological networks by sparse identification of nonlinear dynamics. IEEE Transactions on Molecular, Biological and Multi-Scale Communications. 2016;2(1):52–63.

[2] Kaheman K, Kutz JN, Brunton SL. SINDy-PI: a robust algorithm for parallel implicit sparse identification of nonlinear dynamics. Proc Math Phys Eng Sci. 2020;476(2242):20200279.
